# Dimensional neuroimaging: (Internet) Gaming Disorder symptoms according to the WHO and APA frameworks associate with lower striatal volume

**DOI:** 10.1101/852855

**Authors:** Xinqi Zhou, Renjing Wu, Congcong Liu, Juan Kou, Yuanshu Chen, Halley M. Pontes, Dezhong Yao, Keith M. Kendrick, Benjamin Becker, Christian Montag

## Abstract

Growing concerns about the addictive nature of internet and computer games led to the endorsement of Internet Gaming Disorder (IGD) as an emerging disorder by the American Psychological Association (APA) in 2013 and the recognition of Gaming Disorder (GD) as a new diagnosis by the World Health Organization (WHO) in 2019. While the definition of clear diagnostic criteria for IGD is beneficial for clinicians and those in need of treatment, it may also stigmatize normal behavior. Furthermore, the neurobiological mechanisms underlying IGD symptoms, and whether they resemble those of other addictive disorders remains highly debated. To this end the present study employed a dimensional imaging approach to determine associations between IGD symptom-load according to the APA and WHO diagnostic frameworks and brain structure in a comparably large sample of healthy subjects. It was found that higher symptom-load on psychometric tests assessing the APA and WHO diagnostic frameworks convergently associated with lower volumes of the striatum. The present results suggest for the first time a neurobiological foundation of the proposed diagnostic criteria for IGD according to both diagnostic classification systems, further indicating that the transition from non-disordered to disordered gaming is accompanied by progressive neuroplastic changes in the striatum, thus resembling progressive changes in other addictive disorders.

## Introduction

Growing concerns regarding the potential detrimental effects of excessive Internet gaming on mental health, including the development of addictive patterns of use have led to the inclusion of Internet Gaming Disorder (IGD) in the major classification systems for mental disorders. After reviewing the growing body of research on excessive gaming (Pontes & Griffiths, 2019) the World Health Organization (WHO) formally recognized Gaming Disorder (GD) as an official diagnosis to be included in the next revision of the International Classification of Diseases (ICD, 11^th^ version). Earlier in 2013, the American Psychiatric Association (APA) included IGD as an emerging disorder in the appendix of the DSM-5 (APA, 2013). Although the specific symptoms for disordered gaming differs across the APA and WHO diagnostic frameworks, the proposed diagnostic criteria in both frameworks strongly resemble the diagnostic criteria for substance-based addictions and include preoccupation with gaming, loss of control over gaming and continued use despite negative consequences (details see **Supplemental Materials**). For the categorical diagnosis of IGD a cut-off number of the symptoms must be exhibited over the last 12-months and the symptoms must be severe enough to lead to marked distress and functional impairments in daily life.

By proposing clear guidelines on how to assess and diagnose IGD, the classification systems have provided tremendous help for clinicians, researchers and those in need of treatment for disordered gaming. Furthermore, the recent developments account for growing public concerns about the detrimental effects of excessive gaming on mental health with the Chinese Government even releasing new policies in November 2019 to regulate centrally the time spent on internet gaming by adolescents (https://www.bbc.com/news/world-asia-50315960). On the other hand, the inclusion of IGD as a recognized addictive disorder may also pathologize and stigmatize normal behavior as the neurobiological mechanisms underlying IGD symptoms and their resemblance to changes in other addictive disorders remain highly controversial (Zastrow, 2017). On the clinical and phenomenological level IGD subjects exhibit symptoms that strongly resemble compulsive use and loss of control observed in other addictive disorders, such as substance-based addictions. Converging evidence from animal models and human imaging studies indicate that the transition from volitional to addictive and compulsive drug use in substance-based addictions is accompanied by progressive dysregulations in the motivational circuits of the brain. The striatum lies at the heart of these motivational circuits and plays a critical role in both the initial reinforcing effects of addictive substances as well as the development of compulsive use and progressive loss of control that characterize the transition to substance-based addictions (Koob & Volkow, 2016). Neuroplastic changes in the striatum and associated circuitries engaged in reward processing and habit formation have been demonstrated extensively in animal models of substance-addiction (Everitt & Robbins, 2016). Human neuroimaging studies have also repeatedly reported striatal gray matter alterations in addicted populations (Mackey et al., 2019), with the extent of volumetric reductions being associated with escalating use and clinical symptom severity (Garza-Villarreal et al., 2018; Becker et al., 2015)

Provisional evidence from a growing number of human imaging studies in pathological gambling – an established behavioral addiction – indicates that striatal circuits play an important role in the initial rewarding effects of gambling (Breiter, Aharon, Kahneman, Dale, & Shizgal, 2001) as well as the formation of compulsive gambling during later stages of the disorder (Clark, Boileau, & Zack, 2019). These findings suggest that these circuits may also mediate escalating and ultimately compulsive use in behavioral addictions (see also Brand et al., 2019). Compared to substance-based addictions, brain structural alterations in behavioral addictions such as pathological gambling might be less pronounced, probably due to the lack of neurotoxic effects (Clark et al., 2019), even though a growing body of evidence suggests altered gray matter volume in individuals with IGD, including striatal circuits (Yao et al., 2017). Initial studies in the general population reported associations between the extent of gaming or the level of “internet addiction”-associated symptoms and gray matter volume in reward processing regions, including the striatum (Kuhn et al., 2011; Montag et al., 2017; Zhou et al., 2019), suggesting that progressive volumetric changes in this region may accompany the transition into IGD. With the advent of the Research Domain Criteria approach dimensional imaging approaches have been increasingly employed in normal and pathological populations to map associations between pathology-relevant symptom dimensions and progressive dysregulations in the underlying brain systems (Albaugh et al., 2017; Li et al., 2019).

To determine whether the proposed symptom-level criteria for IGD reflects the hypothesized neurobiological basis of the disorder and if the different symptomatic criteria proposed by the APA and WHO access the same neurobiological markers, the present dimensional neuroimaging study aimed at determining associations between subclinical symptoms of IGD and striatal morphology. To achieve this, the study employed non-parametric voxel-wise regression analyses on high-resolution T1-weighted MRI brain structural images with IGD symptom-load as assessed by diagnostic-system specific scales (IGDS9-SF according to the APA framework; GDT according to the WHO framework) (Pontes & Griffiths, 2015; Pontes et al., 2019) as separate predictors in a comparably large sample of healthy young adults (*n* = 82). Psychopathological symptom load on dimensions such as depression and autism frequently associated with IGD were further controlled for. Given the substantial correlations between IGD and GD scores when assessed with the IGDS9-SF and the GDT (Montag et al., 2019), and the key role of the striatum in substance-based addictions it was expected that higher symptom-load on both disordered gaming scales would be accompanied by stronger alterations in striatal morphology.

## Methods and Materials

### Participants and procedures

For the present study *N* = 256 healthy male individuals from the Chengdu Gene Behavior Project were re-contacted via telephone interviews (*N* = 256, > 18 years old). All subjects underwent structural MRI assessment during the 12 months prior to the assessment of GD symptoms (1.1±0.764 years in the final sample). Following MRI data quality assessments and exclusion of duplicate datasets, subjects were contacted via telephone and underwent an interview to exclude those with current or previous history of psychiatric disorders and current use of medication or psychotropic substances. Among all eligible participants, *N* = 119 agreed to participate in the additional electronic data assessment. The severity of IGD symptoms was assessed in light of the APA and the WHO diagnostic frameworks. Given the existence of high levels of comorbidity between depressive disorder and autism spectrum disorder in disordered gaming (e.g. Li et al., 2019; Xu et al., 2019), the potential confounding effects for these two psychopathologies were controlled in the present study using the BDI-II and the ASQ (Baron-Cohen et al., 2001; Beck et al., 1996). Subjects meeting the clinical cut-off criteria (ASQ > 30; BDI-II > 28) were excluded and symptom severities in both dimensions were additionally included as covariates in the analyses. For the final sample, the following exclusion criteria were applied 1) history or current psychiatric disorders according to DSM-5 (validated by structured clinical interviews at the time of MRI assessment), 2) pathological levels of depressive (BDI >=28) and autistic (ASQ >= 30) symptoms, 3) history of or current medical disorder, including neurological and endocrinological disorders, 4) current or regular use of medication and other psychoactive substances, and 5) left-handedness. Of the eligible participants a total of *n* = 82 individuals (age at MRI assessment = 21.8±2.16 years) provided complete self-report and MRI data and were thus included in the subsequent voxel-based morphometry (VBM) analysis (exclusion of subjects detailed in **Fig.S1**). The study and its procedures had full approval by the local ethics committee and adhered to the most recent version of the Declaration of Helsinki and all participants were required to provided informed consent.

### Questionnaires

In the present study Chinese versions of the IGDS9-SF (Pontes & Griffiths, 2015) and the GDT (Pontes et al., 2019) were administered. The IGDS9-SF assesses IGD according to the APA framework using nine items answered on a five-point Likert scale (from 1 = never to 5 = very often). Moreover, the GDT consists of four items assessing GD according to the WHO framework on a five-point Likert scale (from 1 = never to 5 = very often). Answering with *often* or *very often* on the two tests indicates endorsement of the corresponding diagnostic criterion. Consequently, fulfilling five out of nine criteria on the IGDS9-SF and fulfilling all four criteria on the GDT indicates disordered gaming. Internal consistencies were satisfying for both tests (IGDS9-SF α = .845 and GDT α = .912) in the present sample.

### MRI data acquisition

The data was acquired on a 3.0 Tesla GE MR750 system (General Electric Medical Systems, Milwaukee, WI, USA). T1-weighted high-resolution anatomical images were acquired with a spoiled gradient echo pulse sequence, repetition time (TR) = 6 ms, echo time (TE) = minimum, flip angle = 9°, field of view (FOV) = 256 × 256 mm, acquisition matrix = 256 × 256, thickness = 1 mm, number of slice = 156.

### VBM preprocessing

VBM analysis allows the voxel-wise estimation of the local gray matter volume (Ashburner & Friston, 2005). Structural MRI data were preprocessed with CAT12 implementing the computational anatomy approach (http://dbm.neuro.uni-jena.de/cat) based on SPM12 (Welcome Department of Cognitive Neurology, London, UK, http://www.fil.ion.ucl.ac.uk/spm/software/spm12) running on MATLAB Version 8.3 (Math Works Inc., Natick, MA). The standard VBM preprocessing protocols of CAT12 (outlined in the CAT12 manual) were employed. Briefly, the T1-weight images were bias-corrected, segmented into gray matter (GM), white matter (WM), and cerebrospinal fluid (CSF) and spatially normalized to the standard Montreal Neurological Institute (MNI) space using the East Asian template. GM images were smoothed with a Gaussian kernel of 8 mm full-width at half maximum (FWHM) for subsequent statistical analysis and the total intracranial volume (TIV) was estimated to correct for individual differences in brain size. Default parameters were applied unless indicated otherwise. After preprocessing all images passed visual inspection artefacts and an automated quality protocol. Mean Image Quality Rating (IQR) of the final data was B-(84.37%), indicating excellent image quality in the present sample.

### Statistical analyses

Multiple linear regression models were employed in the Statistical non-Parametric Mapping toolbox (SnPM13, http://www.warwick.ac.uk/snpm) based on 10,000 random permutations, taking GM maps as dependent variable and IGDS9-SF and GDT scores respectively as independent variable (similar approach see Li et al., 2019). Although false positive rates in VBM are not strongly influenced by sample size, a non-parametric estimation has been demonstrated as robust against non-normal distribution of the underlying data and might be more robust independent of sample size (Scarpazza, Tognin, Frisciata, Sartori, & Mechelli, 2015; Silver, Montana, Nichols, & Alzheimer’s Disease Neuroimaging, 2011).

For both multiple linear regression models, age, education level, BDI, ASQ, and total intracranial volume (TIV) were additionally included as covariates. In line with the regional-specific a priori hypothesis on striatal associations, voxel-wise analyses were restricted to the striatum (Fig.1C) encompassing both bilateral ventral and dorsal striatal subregions (description of the masks see Zhou et al., 2019). Within the striatal mask a voxel level threshold of *p* < .05 with FWE multiple comparison correction adjusted for the search volume was applied. To further control for the potential effects of time between the brain structural assessments and the assessment of IGD symptoms the time between assessments was controlled for in additional control analyses. Associations between both symptom-scales with striatal volume remained stable (see **Supplementary Materials**).

**Figure 1:**
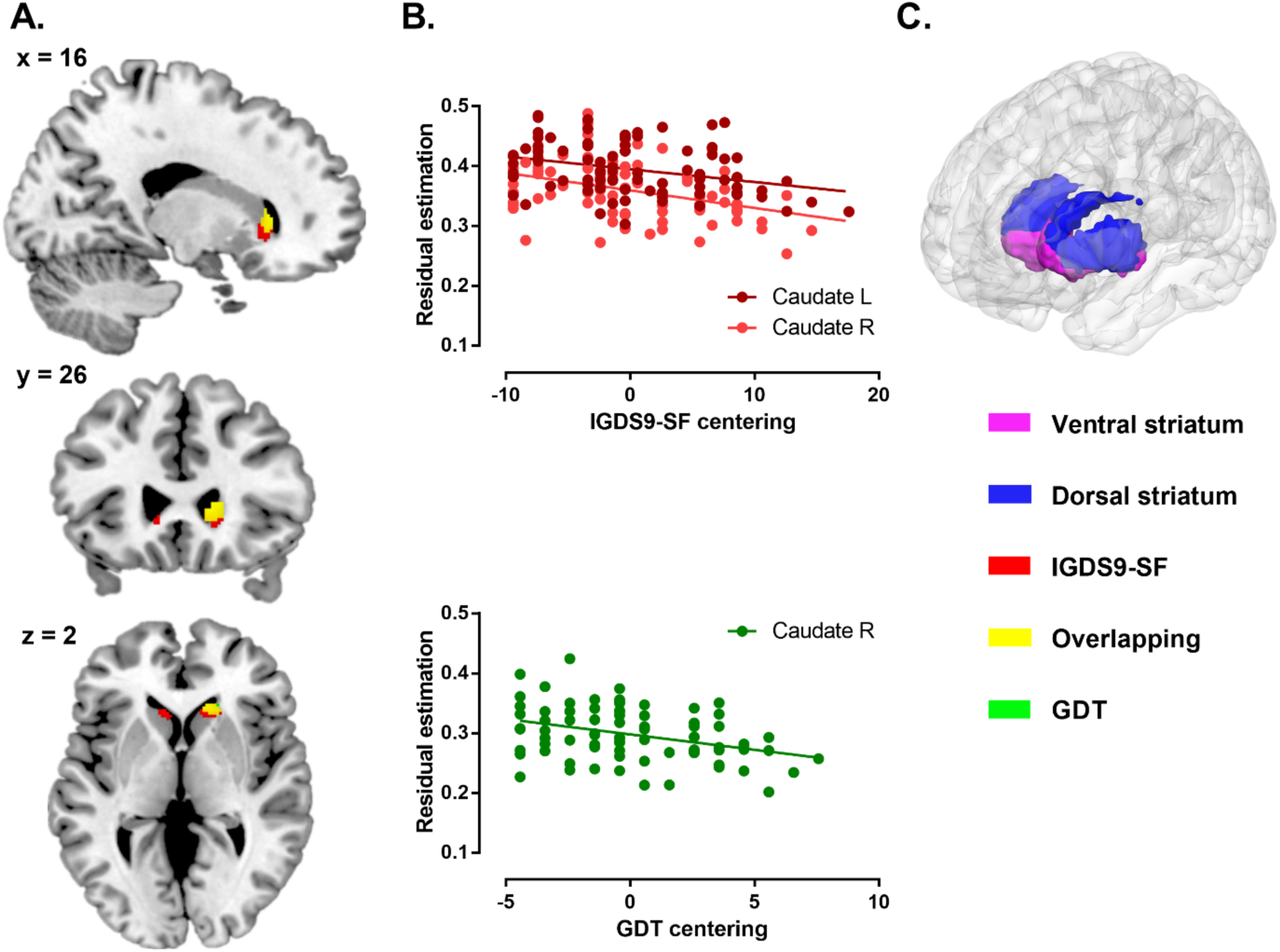
Associations between grey matter volume and IGDS9-SF and GDT. A) Findings were significant at voxel-level pFWE < .05 within the striatum (IGDS9-SF = red, GDT = green, and overlapping = yellow), threshold at k > 30 for displaying purpose) B) The top and bottom panel display extracted volumes from the significant regions and associations with symptom severity C) Ventral and dorsal striatal volumes used in the present study (note that for multiple comparison correction a single mask encompassing the entire bilateral striatum was employed). In general, the IGDS9-SF related structural alteration were more extensive than for the GDT, and encompassed the GDT associated structural alteration.

## Results

### Participants

Categorial analysis of the symptom-level data revealed that in terms of disordered gaming, three subjects fulfilled the diagnostic criteria on the APA framework (IGDS9-SF) and one on the WHO framework (GDT). Dimensional analysis of the symptom level data further revealed that the present sample reported a mean (± SD) symptom severity of = 18.4±6.53 for the IGDS9-SF and 8.43±3.07 for the GDT. Moreover, the mean trait score for depression was 6.26±6.98 (the Beck Depression Inventory, BDI-II, Beck, Steer, & Brown, 1996), and for autism was 13.7±3.77 (the Adult Autism Spectrum Quotient, ASQ, Baron-Cohen, Wheelwright, Skinner, Martin, & Clubley, 2001). Mean total intracranial volume was 1563±107, and education level ranged from high school graduate to master’s degree student.

### Associations between (internet) gaming disorder symptom load and striatal grey matter volumes across the APA and WHO diagnostic frameworks

Higher IGDS9-SF symptom severity was significantly associated with lower gray matter volume in the bilateral caudate (pFWE < .05), whereas higher GDT symptom severity was associated with lower gray matter volume in the right caudate (pFWE < .05) (Table 1, Fig.1 A & B). Given the distinct contributions of the ventral and dorsal striatum to addictive disorders (Zhou et al., 2019) and behavioral addictions (Brand et al., 2019), the significant effects were further mapped and overlapped (Fig.1A), revealing that the IGDS9-SF and GDT shared 114 voxels in caudate volume. Additionally, after resampling the ventral and dorsal striatum masks employed in a previous study conducted by the team on ventral and dorsal striatal contributions to addiction (Zhou et al., 2019) (resampled to match the dimensions of the present structural images), observed clusters for the IGDS9-SF revealed 244 voxels mapping to the ventral striatum and 88 voxels to the dorsal striatum. For the GDT, 53 voxels mapped onto the ventral striatum and 65 voxels on the dorsal striatum.

**Table 1.** Significant negative associations between gray matter volume and the IGDS9-SF and GDT respectively

| Region | *k* (Cluster Size) | x | y | z | *t* Value |
| --- | --- | --- | --- | --- | --- |
| IGDS9-SF, negative associations |  |  |  |  |  |
| Caudate R | 207 | 14 | 26 | 3 | 4.23 |
|  |  | 23 | 24 | 9 | 3.87 |
| Caudate L | 103 | -11 | 24 | -2 | 4.11 |
|  |  | -6 | 21 | 5 | 3.87 |
| Caudate L | 22 | -18 | 21 | 12 | 4.04 |
| GDT, negative associations |  |  |  |  |  |
| Caudate R | 118 | 15 | 27 | 5 | 4.32 |

R = right, L = left.

## Discussion

The present study employed a dimensional imaging approach to determine the neurobiological basis of the recently proposed criteria for IGD according to the WHO and APA diagnostic frameworks. In line with our hypothesis, stronger symptom-load in both diagnostic frameworks was associated with lower striatal gray matter volume, primarily the caudate region. The caudate bridges the ventral and dorsal striatum and is strongly engaged in several addiction-relevant functional domains, with ventral regions being engaged in the initial formation of reward-related salience and associative learning processes (Anderson et al., 2017) whereas more dorsal parts mediate habit formation (Everitt & Robbins, 2016). Exaggerated neural cue-reactivity in the ventral striatum has not only been consistently observed in substance-based addictions but also in non-addicted substance users (Vollstadt-Klein et al., 2010; Zhou et al., 2019). Stronger engagement in reward-related Internet behavior has been associated with lower ventral striatal volume in healthy samples (Montag et al., 2017; Montag et al., 2018), suggesting that neural changes in this region may already occur during early stages of the addictive process. In line with these findings, IGD symptom-load on both scales was consistently associated with lower gray matter volume in this region. This finding provides the first demonstration of a neurobiological correlate of the proposed IGD criteria in the classification systems and further suggests that development of IGD is accompanied by progressive neurobiological changes in brain motivational systems that critically mediate the development of established addictive disorders, including substance-based disorders and pathological gambling (Clark et al., 2019; Koob & Volkow, 2016).

Although the diagnostic criteria of both classification systems are strongly aligned with the respective core symptomatic criteria for substance-based disorders, the specific symptom criteria differ slightly (details **Supplementary Materials**). For scientists and practitioners alike a key issue thus relates to diagnosing essentially the same condition using the two (distinct) diagnostic frameworks (APA vs. WHO). One issue is certainly whether estimation biases can occur when ascertaining prevalence rates of IGD in large-scale research (Montag et al., 2019), whereas a second issue relates to potential neurobiological discrepancies between the two diagnostic frameworks. Examining the overlap between the classification system-specific striatal regions in the present study revealed that symptom-load according to the APA framework captures considerably larger striatal regions, particularly ventral striatal regions. Ventral regions engaged in reward and salience processing are considered to promote reward-driven and pleasure-oriented behavior during early stages of drug use, whereas dorsal regions engaged in habit formation may promote the development of compulsive behavior during later stages of the addictive process (Everitt & Robbins, 2016). The discrepancies in the neurobiological associations may point to a higher sensitivity of the APA framework to detect early stages of disordered gaming. The neural findings align with the higher prevalence of IGD according to APA as compared to WHO criteria in the present sample as well as in a recent large-scale study employing identical scales (Montag et al., 2019). Taken together, these findings may suggest that the WHO framework represents a stricter and therefore, more conservative approach to diagnosing disordered gaming in comparison to the APA framework.

In summary, the present study provides a first demonstration of a neurobiological basis for the proposed IGD criteria in accordance with both current diagnostic systems as well as subtle brain-related differences which are diagnostic framework dependent. The findings suggest that the proposed criteria reflect progressive dysregulations in the motivational systems of the brain. One potential limitation in the present study is that the co-linearity between the IGDS9-SF and GDT does not allow a direct comparison of the imaging results obtained, warranting caution when drawing conclusions in regard to the neural differences between the two psychometric tests. Finally, differences between the diagnostic frameworks warrant further careful consideration when integrating the already existing large body of literature on disordered gaming with future studies employing the WHO framework.

## Disclosure

The authors report no conflict of interest.

