## Supplemental_Material for "Dimensional neuroimaging: (Internet) Gaming Disorder symptoms according to the WHO and APA frameworks associate with lower striatal volume"

### Supplementary Materials

#### Supplemental description of symptoms between two diagnostic frameworks

According to DSM-5 (APA framework) five out of nine symptoms have to be fulfilled over the last twelve months to diagnose IGD. These symptoms are as follows: preoccupation with gaming, withdrawal symptoms when gaming is not possible, development of tolerance, problems in cutting down game play, loss of interest in other daily life activities due to gaming, continuing game play although problems arise in one's own life, lying about one's own actual game play, playing games to reduce negative mood and significant impairments such as losing a job or relationship due to gaming.

Interestingly, the World Health Organization (WHO) proposed somewhat different criteria to diagnose Gaming Disorder (6C51; to be found among the parent category of disorders due to addictive behaviors, https://icd.who.int/browse11/l-m/en#/http://id.who.int/icd/entity/1448597234). In detail, the following criteria all need to be observed to speak of Gaming Disorder. These are: loss of control over gaming, increasing priority of gaming compared to other everyday life activities, continuing gaming despite of negative consequences and significant impairments in private and/or business lives. These symptoms should be visible over the time course of at least one year. Please note the ICD-11 in WHO framework also distinguishes between online and offline Gaming Disorder, whereas DSM-5 only speaks of Internet Gaming Disorder.

#### Supplemental results

*Associations between (internet) gaming disorder symptom load and striatal grey matter volumes across the APA and WHO diagnostic frameworks: controlled for time difference between measurements of questionnaires and T1 data*

Same statistical analysis pipeline was conducted with time difference as additional covariate in multiple regression model. Findings revealed that higher IGDS9-SF symptom severity was significantly associated with lower gray matter volume in the bilateral caudate ($p_{FWE} < .05$, right caudate peak at [12, 26, 2], k = 205, left peak at [-9, 23, 0] and [-18, 21, 12], k = 128 and 15), whereas higher GDT symptom severity was associated with lower gray matter volume in the right caudate ($p_{FWE} < .05$, peak at [15, 27, 5], k = 76).

**Supplemental figure**

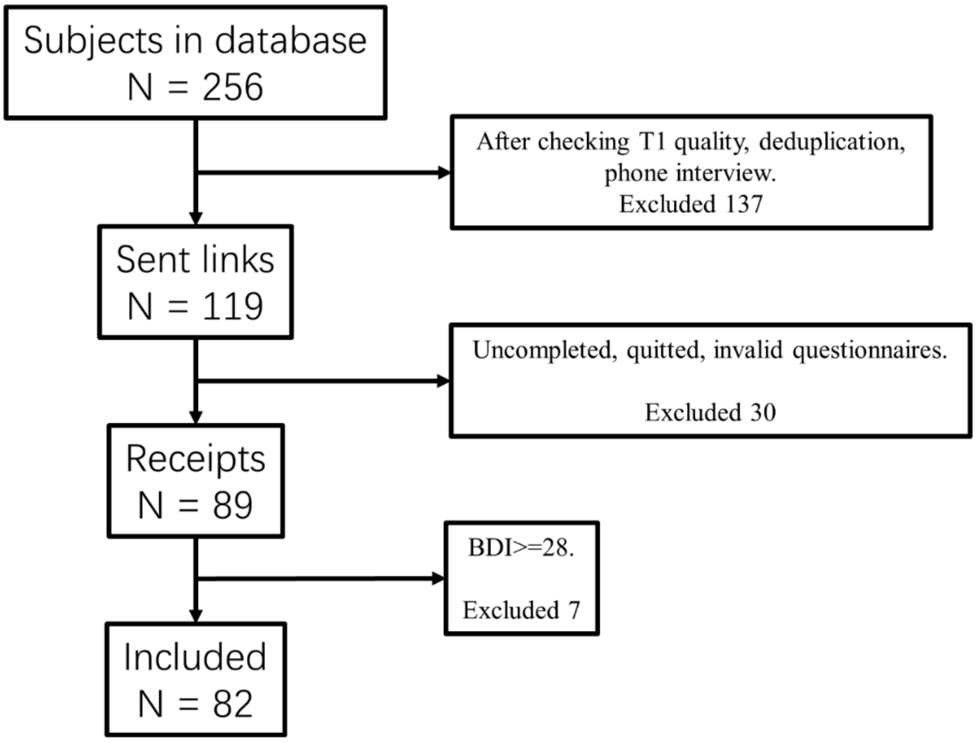


Fig.S1. Flow diagram displaying screening and exclusion of participants
